# HERVs as building blocks of RNA regulatory architecture in the human genome

**DOI:** 10.64898/2026.04.29.721355

**Authors:** Tomàs Montserrat-Ayuso, Aurora Pujol, Anna Esteve-Codina

## Abstract

Human endogenous retroviruses (HERVs) comprise nearly 8% of the human genome and have contributed extensively to gene regulatory evolution. However, their roles in RNA-centered regulatory processes remain poorly characterized. Here, we present a genome-wide annotation of RNA regulatory features embedded within HERV internal regions and long terminal repeats (LTRs), revealing that HERV sequences act as pervasive components of the human transcriptome. Systematic analysis of RNA-binding protein (RBP) motifs uncovers structured, family-specific regulatory architectures, with distinct RBP signatures distinguishing major HERV subfamilies. Notably, HERVH elements are enriched for RBPs associated with developmentally regulated RNA processing, whereas HERVK (HML-2) elements preferentially harbor motifs linked to canonical splicing and mRNA maturation. Integration with gene annotations reveals widespread incorporation of HERV sequences into transcript structures, including more than 4,000 long non-coding RNAs. Conserved retroviral protein domains within predicted open reading frames are strongly enriched in terminal exons and 3′ untranslated regions, consistent with potential micropeptide-encoding capacity. In addition, we identify a subclass of lncRNAs largely composed of HERV sequence, indicating that endogenous retroviral loci have been extensively captured within annotated transcripts. Finally, we detect more than 6,500 antisense LTR insertions in transcript termini, defining widespread SPARCS-like (stimulated 3 prime antisense retroviral coding sequences) configurations with potential for double-stranded RNA formation and preferential association with immune-related genes. Together, these results establish HERV sequences as a pervasive layer of RNA regulatory potential embedded within human transcripts, highlighting previously underappreciated roles in post-transcriptional gene regulation.

## Introduction

Transposable elements (TEs) account for roughly 45% of the human genome^1^ and are increasingly recognized as important contributors to gene regulation^2^. Human endogenous retroviruses (HERVs), a major class of TEs derived from ancient germline infections by exogenous retroviruses, comprise approximately 8% of the genome and represent a particularly rich source of regulatory sequences, including long terminal repeats (LTRs) and internal regions with distinct functional properties^3^. HERV-derived LTRs have been extensively co-opted as promoters and enhancers that shape gene expression programs across development, cellular differentiation, and disease^4,5^. By contrast, the internal regions of HERVs remain comparatively understudied, despite retaining signatures of their retroviral origin and exhibiting varying degrees of evolutionary conservation, in the sense of HMM profile coverage^6^. Their potential contribution to RNA-level regulatory processes has not been systematically explored.

Following their integration, HERV sequences have undergone extensive mutation, recombination, and deletion, resulting in a spectrum of genomic structures ranging from near-intact proviruses to fragmented internal regions and solo LTRs generated through homologous recombination^7^. Canonical proviral insertions are organized as LTR–internal–LTR structures, in which the internal region contains retroviral coding genes, including *gag*, *pol*, and *env*^3^. Although most HERV loci have lost protein-coding capacity, many internal regions retain recognizable coding remnants and partial structural organization^6^. This residual architecture suggests that, beyond their evolutionary origin, HERV internal regions may preserve sequence features with potential functional relevance.

Beyond their structural persistence, HERV sequences have been repeatedly co-opted at both the regulatory and coding level. At the regulatory level, LTRs function as promoters and enhancers that can rewire transcriptional programs, particularly in developmental and immune contexts^8,9^. At the coding level, a small number of HERV-derived genes have been retained and repurposed, most notably the syncytins—retroviral envelope proteins essential for placental development^10,11^. At the same time, dysregulated HERV activity has been associated with a range of pathological conditions, including cancer^12–14^, neurological disorders^15,16^, and autoimmune diseases^17^. These dual roles highlight the importance of distinguishing background HERV transcription from loci that retain functional regulatory or coding potential.

At the RNA level, HERV-derived sequences have the potential to encode diverse regulatory features. These include binding sites for RNA-binding proteins (RBPs)^18,19^, sequences that influence RNA stability, localization, and translation^20–22^, and motifs capable of acting as competing endogenous RNAs through microRNA sequestration^23,24^. In addition, specific genomic configurations involving HERV-derived sequences may give rise to double-stranded RNA (dsRNA) structures capable of activating innate immune sensing pathways^25^. Importantly, these features are expected to operate within the context of host transcripts, including long noncoding RNAs (lncRNAs) and protein-coding genes ^26^. Despite these possibilities, the extent to which such RNA-associated regulatory features are systematically encoded within HERV internal regions—and whether they exhibit structured patterns across subfamilies and loci—remains unclear.

A systematic characterization of RNA-associated regulatory features within HERV internal regions is currently lacking. In particular, it remains unclear how these sequences contribute to post-transcriptional regulation through RBP interactions and how they are integrated into host transcript architectures such as lncRNAs and protein-coding genes. Specific genomic configurations involving HERV sequences have been shown to generate dsRNA and activate innate immune signaling, as exemplified by SPARCS (stimulated 3 prime antisense retroviral coding sequences), where antisense HERVs located in 3′ UTRs of interferon-stimulated genes produce dsRNA through bidirectional transcription^25^. However, the extent to which similar dsRNA-forming architectures occur more broadly across the genome, and how they relate to HERV sequence composition and transcript context, remains unclear. Moreover, an integrative, genome-wide framework linking these distinct regulatory layers has not been established.

Here, we present a comprehensive, multi-layer analysis of RNA-associated regulatory features across 53,176 HERV internal regions and 560,594 HERV LTRs in the human genome. By integrating RBP binding site annotation, transcript overlap with coding and non-coding genes, conserved retroviral domain mapping, and LTR regulatory architecture, we define a heterogeneous landscape of regulatory potential across HERV subfamilies and individual loci. Our analysis identifies subfamily-specific RBP binding patterns, reveals a strong enrichment of conserved domains in terminal exons and 3′ untranslated regions, characterizes a class of HERV-dominated lncRNAs, and uncovers widespread genomic configurations consistent with SPARCS-like dsRNA-forming architectures. Together, these findings establish HERV-derived sequences as a pervasive and previously underappreciated layer of RNA regulatory potential in the human genome. To facilitate further exploration and reuse, all annotations generated in this study are provided as an openly accessible resource via Zenodo (https://doi.org/10.5281/zenodo.19661036).

## Results

### Global overview of ncRNA-associated regulatory features in HERV internal regions

To systematically characterize the regulatory potential of HERV internal regions at the RNA level, we integrated multiple layers of annotation across the human genome, including RBP motifs, transcript overlap with coding and non-coding genes, and previously defined annotations of conserved retroviral protein domains and LTR regulatory architecture^5,6^.

This integrated framework enabled the joint analysis of sequence-derived regulatory features together with genomic and transcriptomic context, allowing both subfamily-level comparisons and locus-specific interrogation.

At a global level, HERV internal regions exhibited widespread but highly heterogeneous enrichment of RNA-associated regulatory features. Overall, 88.5% (47,076/53,176) of loci contained at least one potential RBP binding site (RBPBS), with a median of six distinct RBPs per locus, indicating that RNA regulatory features are pervasive across the HERV repertoire.

To dissect this heterogeneity, we first examined the landscape of RBPBS enrichment across HERV subfamilies.

### RBPBS landscape reveals family- and locus-specific binding potential

To determine whether the observed potential RBPBS landscape reflects biologically meaningful regulatory signals rather than nonspecific sequence composition, we assessed RBPBS enrichment across HERV subfamilies (Supplemental Table 1). For each RBP, enrichment within a given subfamily was quantified relative to all other subfamilies using odds ratios derived from motif counts, with multiple-testing correction applied across RBPs.

This analysis revealed clear and structured differences in RBPBS enrichment between HERV subfamilies, with the most pronounced contrast observed between HERVH and HERVK (HML-2) (Figure 1A). Rather than a uniform distribution of enriched RBPs, each subfamily was associated with a distinct and internally coherent set of RBPs.

**Figure 1.**
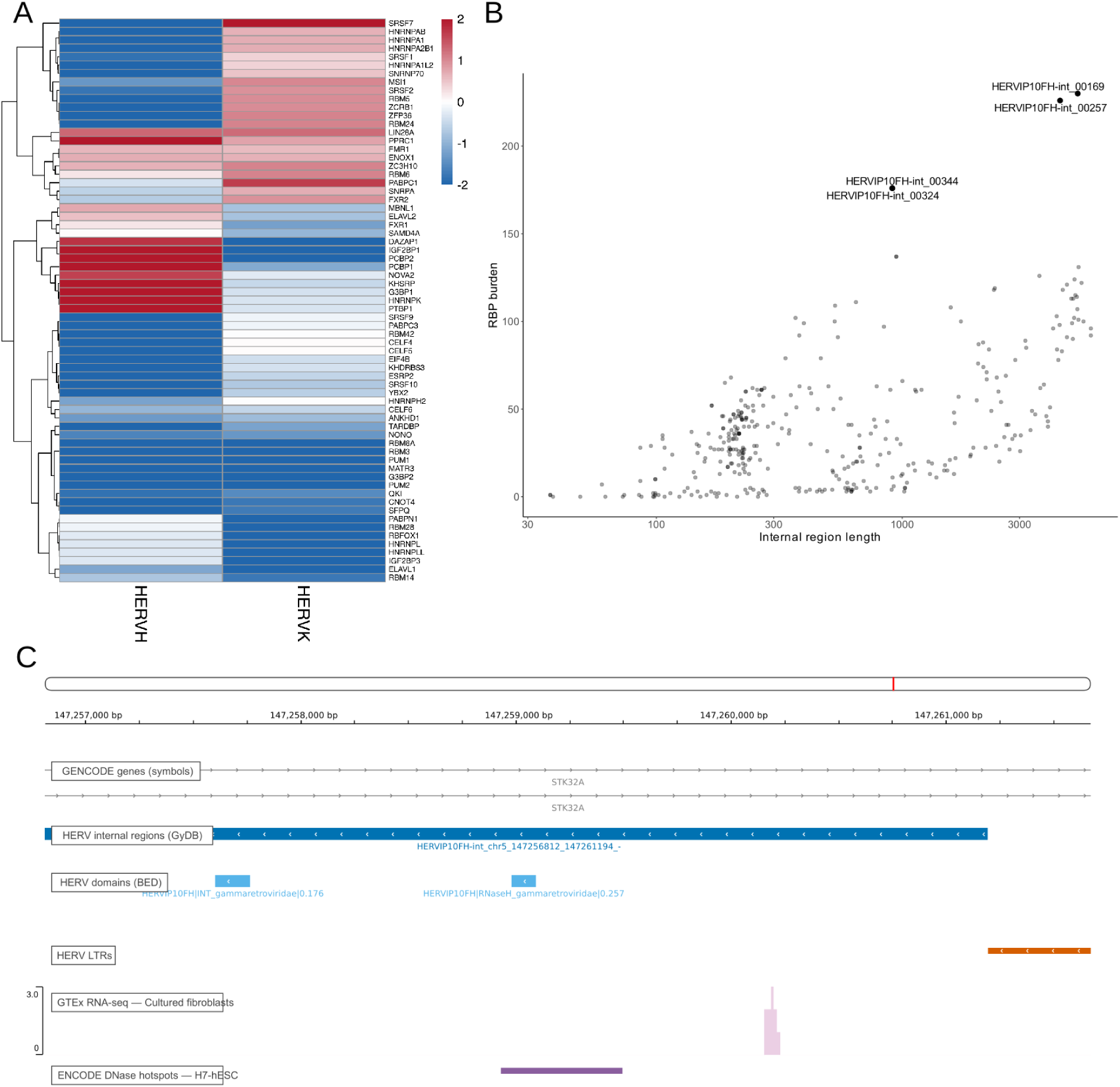
Subfamily- and locus-specific RBP binding potential across HERV internal regions. (A) Heatmap of significant RBP motif enrichment (log2 odds ratio, FDR < 0.05) comparing HERVH and HERVK (HML-2) internal regions, revealing distinct and partially reciprocal RBP binding profiles. (B) Distribution of RBPBS burden as a function of internal region length for HERVIP10FH loci, highlighting extreme outliers with unusually high RBPBS density. (C) Genomic context of a representative outlier locus (chr5:147,256,812–147,261,673) located antisense within an intron of STK32A, showing overlap with HERV internal regions and domains, GTEx RNA-seq signal, and an ENCODE DNase I hypersensitivity hotspot. Visualization generated using the HERVarium web application (https://hervarium.cnag.eu/).

HERVH internal regions were preferentially enriched for motifs corresponding to RBPs involved in developmentally regulated and post-transcriptional RNA control. These included factors linked to pluripotency and early developmental states, such as LIN28A^27^ and IGF2BP1^28^, as well as RBPs implicated in regulated splicing and RNA fate decisions with phenotypic impact, including PTBP1^29^, MBNL1^30^, NOVA2^31^, DAZAP1^32^, HNRNPK^33^, and ELAVL2^34^. Notably, enriched motifs also corresponded to RBPs involved in RNA stability, localization, and translational control, including PCBP1/2^35^, G3BP1^36^, KHSRP^37^, FMR1^38^, and FXR1^39^.

Many of these RBPBS — including PTBP1, MBNL1, HNRNPK, IGF2BP1, and ELAVL2— were enriched in HERVH while being depleted in HERVK (HML-2), revealing a reciprocal association of regulatory splicing and RNA fate factors between the two families.

In contrast, HERVK (HML-2) internal regions were enriched for RBPBS associated with canonical co-transcriptional exon definition and early messenger ribonucleoprotein (mRNP) maturation, including multiple SR proteins (SRSF1, SRSF2, SRSF7) and U1/U2 small nuclear ribonucleoprotein (snRNP) components (SNRPA, SNRNP70)^40^. Alongside these factors, enrichment of several heterogeneous nuclear ribonucleoproteins (hnRNP) proteins (HNRNPA1, HNRNPA2B1, HNRNPAB) suggests additional layers of co-transcriptional splicing modulation^41^. Additional enrichment of RBPBS involved in general mRNA handling, turnover or regulation of translation, such as PABPC1^42^, ZFP36^43^, MSI1^44^, and RBM24^45^, is consistent with HERVK-derived RNAs being preferentially processed through canonical mRNA lifecycle pathways, and potentially subjected to post-transcriptional regulatory mechanisms that limit RNA stability or translation.

Importantly, a limited subset of RBPs associated with pluripotency, including LIN28A, were enriched in both HERVH and HERVK internal regions, consistent with previous reports of HERVK activity in pluripotent contexts^46,47^. Taken together, these results point to partially reciprocal enrichment patterns, with RBPs involved in regulatory and developmentally controlled RNA processing preferentially associated with HERVH, and RBPs linked to constitutive splicing and RNA turnover more frequently associated with HERVK.

### HERVIP10FH as a case study: a candidate co-opted locus embedded in the STK32A gene

To illustrate how subfamily-level analyses can reveal discrete candidate loci with unusually high regulatory potential, we focused on HERVIP10FH as a representative case study. This subfamily comprises a relatively large number of annotated loci in our dataset (n = 353), providing sufficient statistical power to characterize its internal distribution of features while remaining tractable for detailed inspection.

Across HERVIP10FH loci, internal regions displayed a tight and well-defined background distribution of both sequence length and RBPBS burden (median length: 352 bp; median total RBPBS burden: 33; median unique RBPs: 9). Against this background, only a small number of extreme outliers were observed (Figure 1B). Specifically, four loci exhibited more than 150 RBPBS, markedly exceeding the subfamily-wide distribution. Two of these loci are located within the pseudoautosomal regions of the X and Y chromosomes, while the remaining two are embedded within introns of host genes.

Among the intronic outliers, a HERVIP10FH locus located at chr5:147,256,812–147,261,673 stood out as the most extreme example. This locus harbors 226 RBPBS, corresponding to 37 unique RBPs, placing it well beyond the upper tail of the HERVIP10FH distribution.

Genomically, this HERVIP10FH insertion is located antisense within an intron of the gene STK32A. Notably, the element overlaps with intronic RNA-seq signal detected in GTEx cultured fibroblasts, indicating transcriptional activity in a primary human cell context. Independently, the same region overlaps with a DNase I hypersensitivity hotspot from ENCODE, detected in H7 human embryonic stem cells, suggesting an open chromatin configuration in at least one cellular context (Figure 1C). Together, these observations identify this HERVIP10FH insertion as a high-confidence candidate for regulatory co-option, distinguished by extreme RBPBS density, intronic antisense localization, and concordant transcriptional and chromatin accessibility evidence.

### Genomic context of conserved HERV domains reveals preferential retention in terminal exons and 3′ UTRs

To place conserved HERV-derived domains within their transcriptional context, we integrated internal HERV annotations with gene models and regulatory features across the human genome. Conserved domains were defined as those with at least 40% coverage of the corresponding HMM profile. At a global level, 21% of HERV internal regions (11,397 loci) overlapped at least one annotated lncRNA on the same strand, corresponding to 4,074 unique lncRNA genes. In contrast, 6.6% of HERV internal regions (3,514 loci) overlapped protein-coding genes on the same strand, involving 1,521 unique genes. Across all HERV internal regions, intronic overlaps predominated in both gene classes, accounting for ∼6.2% of all loci in protein-coding genes and ∼20.5% in lncRNAs (Figure 2A). Exonic incorporation was comparatively rare in protein-coding genes (∼0.8% of all HERV internal regions, 442 loci), but more frequent in lncRNAs (∼4.6%, 2,467 loci), indicating a relative relaxation of constraints in non-coding transcripts. These categories are not mutually exclusive, as individual HERV internal regions may overlap intronic and exonic regions across different transcript isoforms.

**Figure 2.**
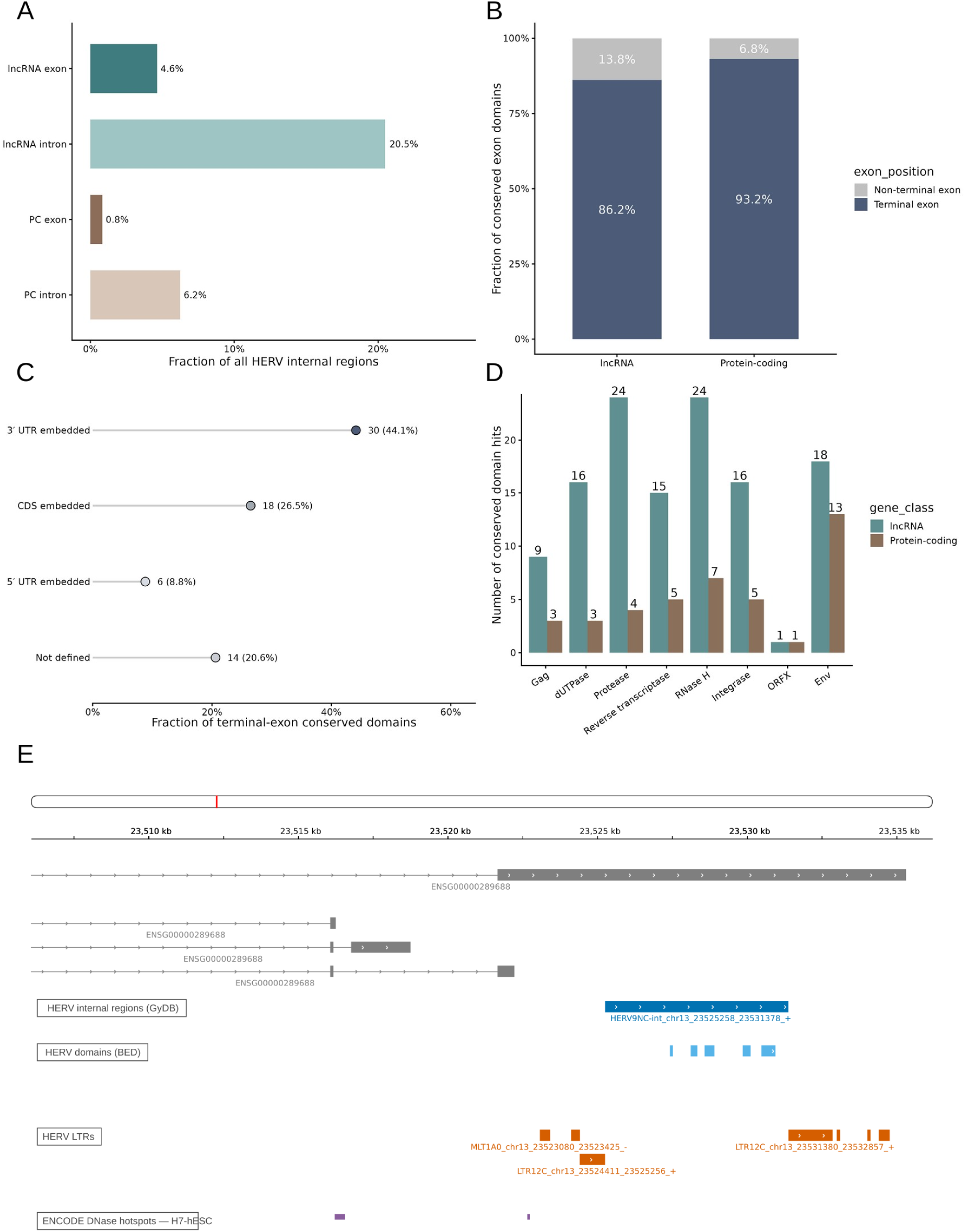
Conserved HERV-derived domains show strong enrichment in terminal exons and 3′ UTRs. (A) Fraction of all HERV internal regions overlapping same-strand lncRNA or protein-coding (PC) introns and exons. (B) Fraction of conserved exon-overlapping HERV domains located in terminal versus non-terminal exons in lncRNA and protein-coding transcripts. (C) Distribution of terminal-exon conserved domains in protein-coding transcripts across 3′ UTR, CDS, 5′ UTR, or transcripts lacking defined CDS/UTR annotation. (D) Domain-class composition of terminal-exon conserved domain hits in lncRNA and protein-coding genes. (E) Genome-browser example of the lncRNA ENSG00000289688, showing a terminal-exon HERV9NC insertion containing multiple conserved HERV-derived domains and flanked by an ENCODE DNase signal. Visualization generated using the HERVarium web application (https://hervarium.cnag.eu/).

We next focused on HERV-derived protein domains exhibiting at least moderate evolutionary conservation (≥40% HMM profile coverage) and overlapping annotated exons in a stringent manner (≥80% of the domain length covered by a single exon). Under these criteria, we identified 264 conserved domains, spanning 90 unique HERV internal regions embedded within 88 distinct genes. Strikingly, these exon-overlapping conserved domains were overwhelmingly enriched in terminal exons, with 88.3% (233/264) located in last exons.

These domains overlapped transcripts from diverse biotypes, including 61 lncRNA genes (188 transcripts) and 24 protein-coding genes (73 transcripts), as well as a small number of TEC (To be Experimentally Confirmed) and processed pseudogene annotations. Notably, 86.2% of lncRNA and 93.2% of protein-coding transcript overlaps involving conserved HERV domains occurred within terminal exons, indicating a strong positional bias of conserved HERV domains toward transcript termini irrespective of gene class. (Figure 2B).

Within protein-coding genes, a minority of last-exon overlaps corresponded to annotated coding sequences (18 transcripts from 8 genes), encompassing well-characterized cases such as ERVW-1 (Syncytin-1) and ERVFRD-1 (Syncytin-2), as well as less explored loci including ERV3-1, where an Env domain is cleanly embedded within the second (terminal) exon of ZNF117. Additional examples included near-complete HERVK (HML-2) insertions with multiple conserved domains retained within exonic contexts.

More frequently, conserved HERV domains were located within 3′ UTRs (Figure 2C). We identified 30 transcripts from 12 protein-coding genes in which conserved domains—including Env, integrase, reverse transcriptase, RNase H, and dUTPase—were embedded within extended 3′ UTRs. Across terminal exons, these domains comprise multiple retroviral classes (Figure 2D). Representative cases included RBM41 and B3GNT7, where highly conserved Env domains occupy long 3′ UTRs, as well as TMEM161B, in which an integrase domain initiates precisely at the end of the canonical coding sequence. These loci frequently coincided with nearby ENCODE candidate cis-regulatory elements (cCREs), either overlapping the HERV insertion itself or flanking the terminal exon, linking conserved HERV-derived coding potential to transcriptionally active gene neighborhoods.

In addition to protein-coding genes, several lncRNAs harbored highly conserved HERV-derived domains embedded within their terminal exons. For example, the lncRNA ENSG00000289688 contains a HERV9NC insertion encoding multiple conserved domains, including protease, RNase H, reverse transcriptase, and Env, with HMM coverage values reaching up to 0.96 (Figure 2E). Notably, ENCODE cCREs overlapped directly with the RNase H and Env domains, situating this locus within a transcriptionally active regulatory context (Supplemental Figure 1A). A similar configuration was observed for ENSG00000285958, where highly conserved RNase H and protease domains (coverages of 0.6 and 1.0, respectively) coincide with multiple overlapping cCREs across the terminal exon (Supplemental Figure 1B). These examples illustrate that the pronounced terminal-exon bias observed genome-wide extends to lncRNAs and is not restricted to protein-coding genes.

Together, these results reveal a pronounced architectural bias whereby evolutionarily conserved HERV protein domains are preferentially retained within last exons and 3′ UTRs. This positional enrichment is observed consistently across both lncRNA and protein-coding genes, indicating that terminal transcript regions represent a recurrent genomic context for the retention of conserved HERV-derived domains.

### Quantitative overlap analysis identifies a class of HERV-dominated lncRNAs HERV internal regions frequently contribute to lncRNA transcript architecture

To assess the contribution of HERV internal regions to annotated lncRNAs, we intersected the catalogue of internal HERV loci with the genomic coordinates of human lncRNA genes. In total, 4,074 lncRNAs were found to overlap at least one HERV internal region, indicating that retroviral sequences are common components of lncRNA loci.

However, the degree of retroviral contribution varied widely across transcripts. To quantify this variation, we calculated the fraction of each lncRNA gene body overlapping HERV internal sequence and classified transcripts according to the extent of HERV contribution.

The vast majority of HERV-associated lncRNAs contained only small retroviral fragments, with 3,043 lncRNA genes harbouring HERV sequence across less than 10% of their gene body (Figure 3A). In contrast, a smaller subset of loci showed substantial retroviral composition. Specifically, 748 lncRNA genes contained between 10–50% HERV sequence, while 139 genes were dominated by HERV sequence (50–80% of the gene body). Notably, 144 lncRNA genes consisted almost entirely of HERV internal sequence (>80% coverage), indicating that these loci largely correspond to retroviral transcriptional units that have been incorporated into the annotated lncRNA repertoire (Figure 3B).

**Figure 3.**
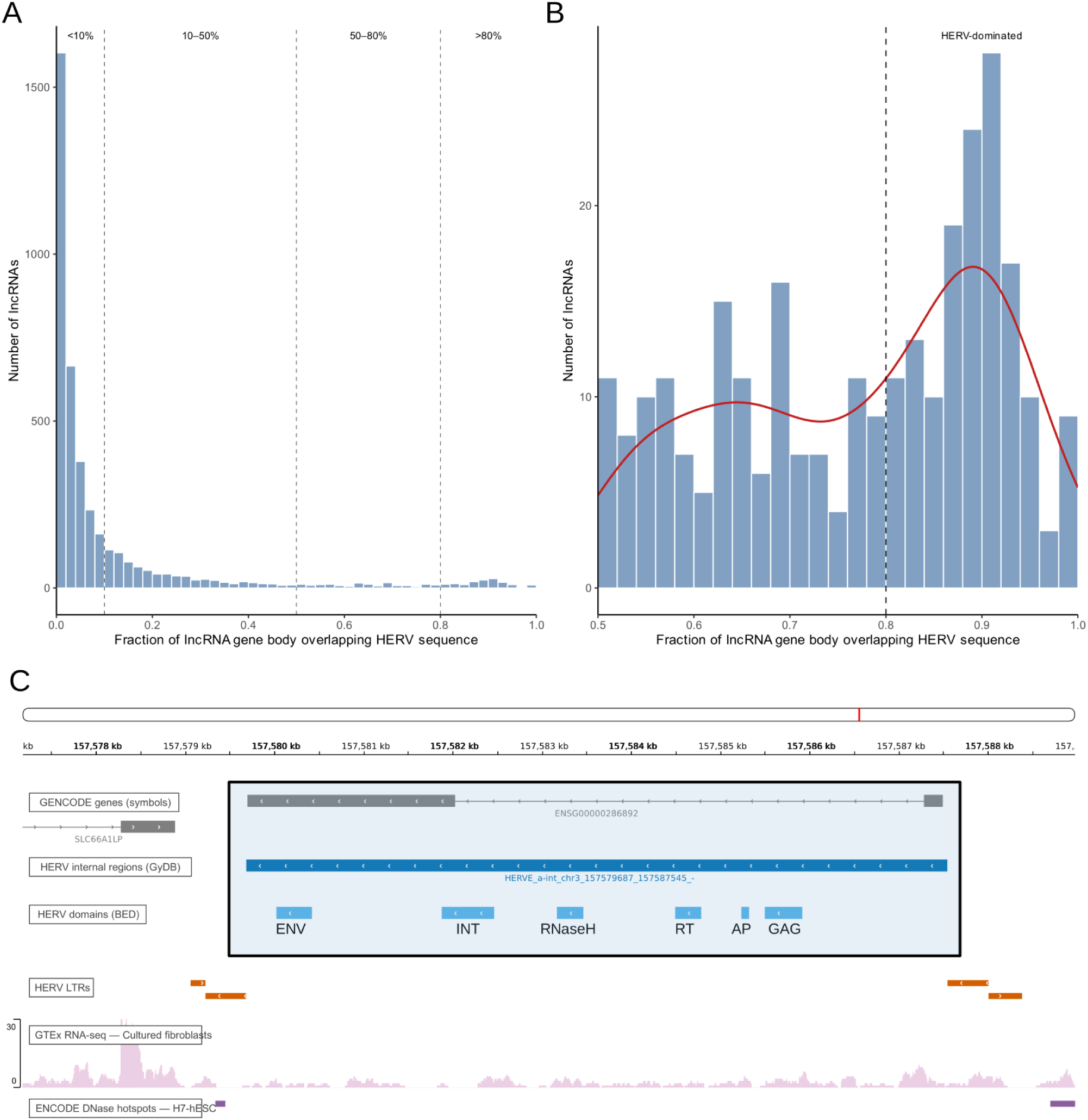
Quantitative contribution of HERV internal regions to lncRNA architecture reveals a subclass of HERV-dominated transcripts. (A) Distribution of the fraction of lncRNA gene bodies overlapping HERV internal sequence. Most lncRNAs contain only limited HERV-derived sequence, with a strong enrichment near low coverage values. Dashed lines indicate classification thresholds used to define HERV contribution categories (<10%, 10–50%, 50–80%, and >80%). (B) Zoomed view of lncRNAs with high HERV contribution (≥50%), highlighting a subset of transcripts with extensive HERV coverage. The dashed line marks the threshold for HERV-dominated lncRNAs (>80% of the gene body). The density curve illustrates enrichment of lncRNAs with high HERV content within this subset. (C) Representative genome-browser view of a HERV-dominated lncRNA locus (ENSG00000286892), in which the entirety of the gene body coincides with a HERV internal region (HERV-E). The locus contains multiple recognizable retroviral protein domains (ENV, integrase, RNase H, reverse transcriptase, protease, and Gag) and is flanked by LTRs, illustrating how endogenous retroviral insertions can form the primary structural and transcriptional unit of annotated lncRNAs. Transcriptional activity is supported by GTEx RNA-seq signal in cultured fibroblasts, and local regulatory potential is indicated by an ENCODE DNase I hypersensitivity signal in human embryonic stem cells. Visualization generated using the HERVarium web application (https://hervarium.cnag.eu/), with highlighting added for clarity.

These HERV-dominated lncRNAs are therefore unlikely to represent host genes that incidentally contain transposable element fragments. Instead, they likely reflect transcription of endogenous retroviral loci that have been captured by genome annotation pipelines as lncRNAs (Figure 3C).

We performed an analogous analysis in protein-coding genes. Although 1,521 protein-coding genes overlapped at least one HERV internal region, extensive HERV contribution was rare. The highest-coverage cases corresponded primarily to known co-opted retroviral genes, including ERVW-1 (syncytin-1), ERVH48-1 (suppressyn), and ERVFRD-1 (syncytin-2), as well as a small number of additional loci annotated with retroviral features (e.g., ENSG00000293569 and ENSG00000293570, annotated as ENV proteins) (Supplemental Table 2). In contrast to lncRNAs, we did not observe a substantial class of protein-coding genes dominated by HERV sequence, consistent with strong selective constraints acting on coding regions.

### Functional annotation of HERV-associated lncRNAs is sparse

To evaluate whether HERV-associated lncRNAs have known biological functions, we integrated curated annotations from the lncRNAWiki database together with literature-derived functional descriptions obtained through automated mining of PubMed titles and abstracts.

Despite the large number of HERV-associated lncRNAs identified, functional annotation was limited. Only 593 of the 4,074 lncRNAs showed any functional description or literature association, while 3,481 transcripts lacked any curated annotation, Gene Ontology (GO) terms^48,49^, or literature evidence, highlighting the largely unexplored nature of this transcript class (Figure 4A).

**Figure 4.**
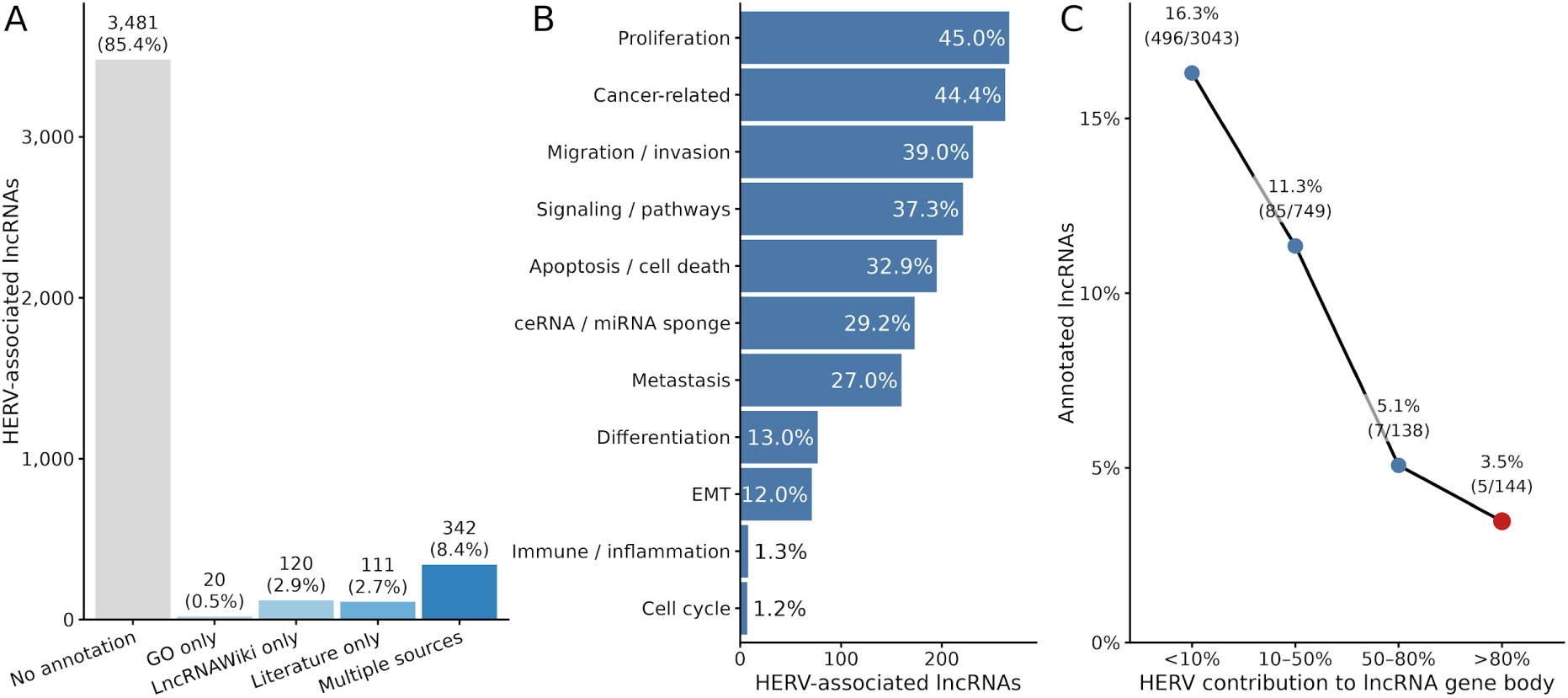
Functional annotation landscape of HERV-associated lncRNAs. (A) Distribution of functional annotation sources across HERV-associated lncRNAs. The majority of transcripts lack any functional annotation (85.4%), while a minority are supported by GO, LncRNAWiki, literature-derived evidence, or multiple sources. (B) Functional categories associated with annotated HERV-associated lncRNAs. Categories were derived by grouping curated and literature-mined annotations into broad, non-mutually exclusive biological processes. The most frequent associations include proliferation, cancer-related processes, migration/invasion, signaling, apoptosis, and ceRNA/miRNA sponge activity. (C) Relationship between HERV contribution and functional annotation. The proportion of annotated lncRNAs decreases progressively with increasing HERV content, reaching near-complete absence among HERV-dominated transcripts (>80% of gene body).

Among the lncRNAs with functional annotation, reported biological associations were dominated by categories related to proliferation, cancer, migration/invasion, signaling, apoptosis, and metastasis. At the mechanistic level, one of the most recurrent associated modes of action was competing endogenous RNA (ceRNA) activity, whereby lncRNAs act as miRNA sponges to modulate post-transcriptional regulation. In total, 173 HERV-associated lncRNAs were linked to ceRNA-related annotations (Figure 4B).

Strikingly, functional information was almost entirely absent for the most HERV-enriched lncRNAs. Of the 144 lncRNAs composed of more than 80% HERV internal sequence, only five genes (3.5%) showed any functional annotation or literature evidence (Figure 4C). Among these, only two genes (CERNA2 and LINC02617) had curated annotation in the lncRNAWiki database.

Together, these results indicate that while HERV sequences are widespread components of lncRNA loci, the biological roles of most HERV-derived transcripts remain largely unexplored.

### Widespread antisense HERV insertions define SPARCS-like transcript architectures with dsRNA-forming potential

Antisense HERV insertions within transcript termini have the potential to generate complementary RNA molecules capable of forming dsRNA structures that activate innate immune sensing pathways. A subset of such elements, termed SPARCS (stimulated 3′ prime antisense retroviral coding sequences), has been described in cancer cells as interferon-inducible HERVs located in the 3′ UTRs of specific genes, where bidirectional transcription generates dsRNA and triggers innate immune signaling^25^.

To investigate whether similar genomic configurations are more broadly present in the human transcriptome, we systematically identified antisense LTR insertions located within the 3′ regions of annotated transcripts. Because our analysis is based on genomic configuration rather than transcriptional activation or functional validation, we refer to these elements as SPARCS-like candidates (Figure 5A).

**Figure 5.**
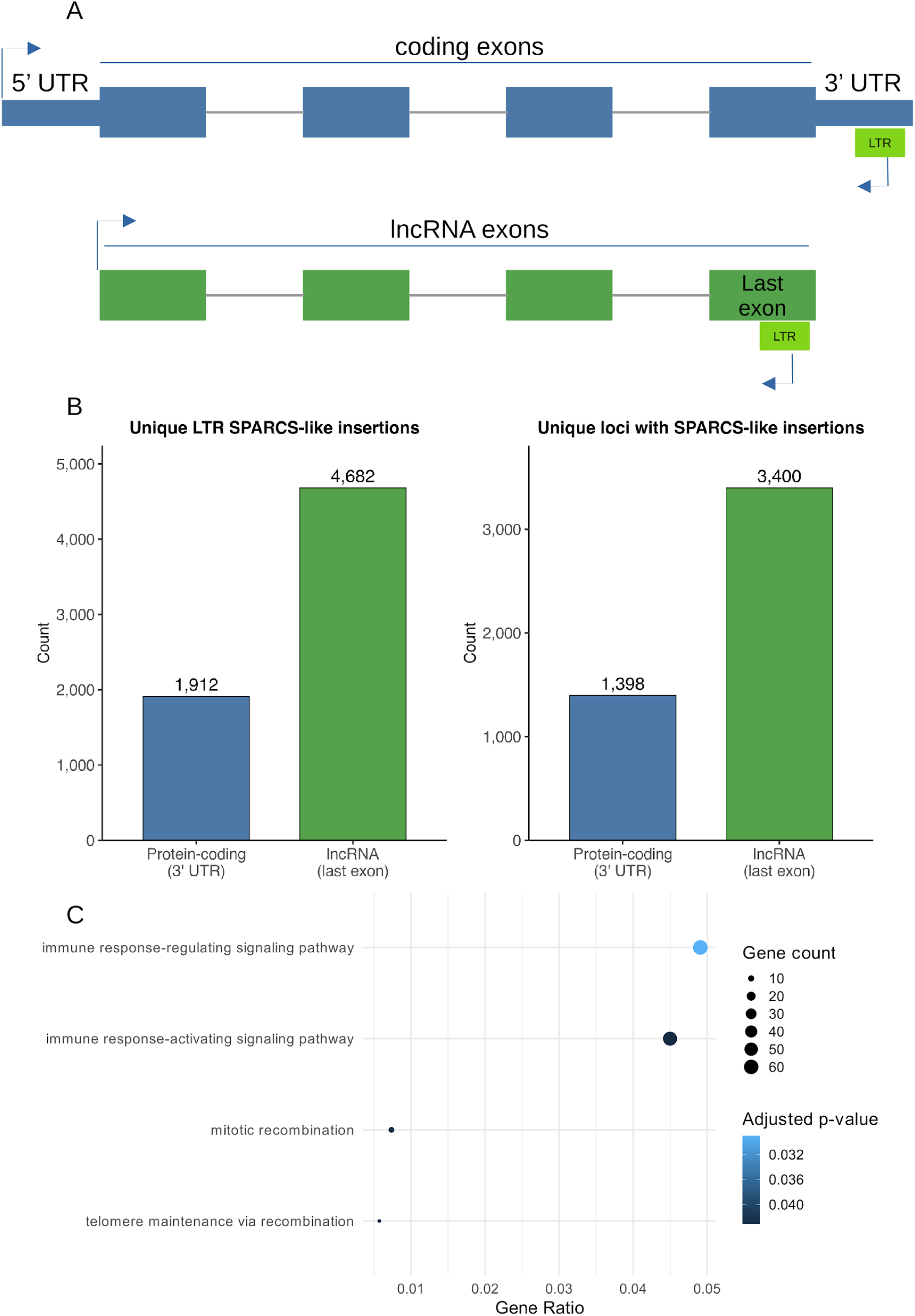
Widespread antisense LTR insertions define SPARCS-like transcript architectures. (A) Schematic representation of SPARCS-like configurations. In protein-coding genes, antisense LTR insertions are located within 3′ UTRs, whereas in lncRNAs they are frequently embedded within the last exon. In both cases, antisense orientation relative to the host transcript creates the potential for bidirectional transcription and dsRNA formation. (B) Genome-wide quantification of SPARCS-like candidates. Left: number of unique antisense LTR insertions located within transcript termini. Right: number of affected genes or loci. Antisense LTR insertions are more frequent in lncRNA last exons than in protein-coding 3′ UTRs, affecting a larger number of loci. (C) GO enrichment analysis of protein-coding genes harboring SPARCS-like insertions. Dot size represents gene count and color indicates adjusted p-value. Enriched terms are predominantly associated with immune signaling pathways.

Across the human genome, we identified 1,912 antisense LTR insertions located within the 3′ UTRs of protein-coding genes, affecting 1,398 genes. In addition, we detected 4,682 antisense LTR insertions located within the last exons of lncRNAs, corresponding to 3,400 distinct lncRNA loci (Figure 5B). In total, these analyses identify more than 6,500 antisense LTR insertions located within transcript termini, indicating that the genomic potential for SPARCS-like configurations is widespread in the human transcriptome.

Analysis of the genomic context of SPARCS-like candidates revealed that the majority occur within non-coding transcripts. Specifically, 60% of affected genes correspond to lncRNAs, whereas 34% correspond to protein-coding genes, with the remaining 6% distributed across other transcript categories, including miRNAs, miscRNAs, processed pseudogenes, rRNA pseudogenes, snoRNAs, scaRNAs, and TEC transcripts.

These results indicate that antisense LTR insertions with the potential to generate dsRNA structures are particularly frequent within non-coding transcript contexts—a setting not explored in previous studies of SPARCS—although a substantial fraction also occurs within the 3′ regions of protein-coding genes.

GO enrichment analysis of protein-coding genes harboring SPARCS-like candidates revealed significant overrepresentation of immune-related biological processes, specifically immune response–regulating signaling pathways and immune response–activating signaling pathways (Figure 5C).

Consistent with this observation, enrichment analysis using immune-related gene sets from the Molecular Signatures Database (MSigDB) (Liberzon et al. 2011) identified three significantly enriched immune signatures: GSE42021_TREG_PLN_VS_CD24INT_TREG_THYMUS_DN, GSE23925_LIGHT_ZONE_VS_DARK_ZONE_BCELL_DN, and GSE42021_CD24HI_VS_CD24LOW_TCONV_THYMUS_DN (Supplemental Table 3). Several genes harboring SPARCS-like insertions are known components of innate immune sensing and interferon signaling pathways. These include interferon-stimulated genes such as OAS1, OAS3, OASL, DDX60, EIF2AK2 (PKR), and GBP5; genes involved in pattern-recognition receptor signaling including TLR1, TLR3, TLR5, TLR8, and MAVS; immune signaling regulators such as NLRP1, BIRC3, CARD8, and TIFA; and adaptive immune signaling components including PTPRC (CD45), LCP2 (SLP-76), CD8B, and CCR7.

Together, these results indicate that SPARCS-like antisense LTR insertions preferentially occur in genes associated with immune signaling pathways.

To determine whether SPARCS-like LTRs exhibit regulatory signatures associated with interferon responses, we examined the enrichment of interferon-responsive transcription factor binding motifs, specifically STAT1, STAT1::STAT2, and members of the IRF family^50–52^. SPARCS-like LTRs were neither enriched nor depleted for these motifs, and none were associated with clustered STAT1/IRF motif architectures indicative of strong interferon-responsive regulation.

Motif analysis further revealed a regulatory landscape distinct from that observed for LTRs acting as promoters. In particular, motifs associated with developmental transcription factors—especially members of the HOX family and other patterning regulators—were significantly depleted in SPARCS-like LTRs located within lncRNA transcripts. GO analysis of these depleted transcription factors highlighted biological processes related to regionalization and pattern specification (Supplemental Table 4 and Supplemental Figure 2).

These observations indicate that SPARCS-like LTRs exhibit a regulatory landscape distinct from promoter-associated LTRs, which are broadly enriched for transcription factors involved in developmental programs^4,5^.

We next examined whether SPARCS-like LTRs display features indicative of increased sequence preservation. Using previously reconstructed canonical U3–R–U5 annotations^5^, we did not observe increased structural integrity relative to other genomic LTRs, with similar proportions of loci exhibiting plausibly reconstructed structures. These results indicate that SPARCS-like configurations are not preferentially associated with structurally intact LTRs.

Together, these findings support a model in which antisense HERV insertions define a widespread transcript architecture that may facilitate dsRNA formation independently of classical interferon-responsive regulatory signatures or enhanced LTR structural preservation.

## Discussion

HERVs have traditionally been studied as sources of DNA-level regulatory elements, with particular emphasis on LTRs acting as promoters and enhancers that shape host transcriptional programs. In this study, we extend this paradigm by demonstrating that HERV sequences also encode a pervasive and structured layer of RNA-centered regulatory potential, embedded within the human transcriptome. Rather than functioning solely as isolated regulatory elements, HERV-derived sequences are extensively incorporated into host transcripts, where they may contribute diverse features that operate at the post-transcriptional level.

Our integrative analysis reveals that this RNA regulatory landscape is inherently multi-layered, arising from the convergence of several distinct but interconnected properties. First, HERV internal regions harbor widespread and non-random distributions of RBPBS, defining subfamily-specific motif-based RBP interaction profiles that suggest structured engagement with cellular RNA regulatory networks. Second, HERV sequences are broadly embedded within transcript architectures, including both long ncRNAs and protein-coding genes, where they contribute to intronic, exonic, and particularly terminal exon contexts. Third, conserved retroviral protein domains are preferentially retained within transcript termini, especially within last exons and 3′ untranslated regions, indicating that remnants of ancestral coding capacity persist in positions compatible with RNA-level regulation and potential micropeptide production. Finally, specific genomic configurations involving antisense LTR insertions give rise to widespread SPARCS-like architectures with the potential to generate dsRNA, linking HERV sequence organization to innate immune sensing pathways.

Taken together, these observations support a unifying model in which HERV sequences provide embedded RNA regulatory modules at the level of sequence features and transcript context, suggesting roles beyond passive genomic relics or isolated cis-regulatory elements. Within this framework, individual HERV loci contribute combinatorial regulatory features—including RBP interaction potential, structural integration into transcripts, residual coding capacity, and dsRNA-forming configurations—that collectively shape RNA behavior in a context-dependent manner. This perspective shifts the interpretation of HERVs from discrete elements with singular functions to components of a distributed regulatory system operating across multiple layers of RNA biology, spanning RNA–protein interactions, transcript architecture, translational potential, and RNA structural organization.

A central insight emerging from this work is that the RNA regulatory potential encoded by HERV sequences is not uniformly distributed across the genome, but instead exhibits subfamily-specific specialization at the level of RBP-binding landscapes. The structured enrichment of RBPBS across HERV internal regions indicates that different HERV subfamilies engage distinct components of the host RNA regulatory machinery.

The clearest contrast is observed between HERVH and HERVK (HML-2), which display partially reciprocal RBP enrichment profiles. HERVH elements are preferentially associated with RBPs involved in developmentally regulated RNA processing, including factors linked to pluripotency, alternative splicing, and RNA fate control, consistent with their activity in early developmental contexts^53,54^. In contrast, HERVK elements are enriched for RBPs associated with canonical co-transcriptional splicing, exon definition, and mRNA maturation, suggesting tighter integration into constitutive RNA processing and surveillance pathways.

These observations support a model in which distinct HERV subfamilies encode different post-transcriptional regulatory programs shaped by both their evolutionary origins and their transcriptional environments. Differences in sequence composition and evolutionary age likely contribute to these patterns, with younger families such as HERVK^7^ retaining features resembling exogenous retroviral transcripts, whereas older families such as HERVH appear more extensively co-opted into specialized regulatory networks^55–57^. Within this framework, HERV sequences can be viewed as modular RNA regulatory cassettes with subfamily-specific biases further modulated at the locus level by genomic context and sequence divergence.

Beyond encoding sequence-level regulatory features, HERV insertions are extensively integrated into the structural organization of host transcripts, contributing to both coding and non-coding RNA architectures. This widespread incorporation indicates that HERVs function not only as sources of regulatory motifs, but also as structural modules that shape how transcripts are assembled and regulated^58^.

Our analyses reveal that HERV sequences are broadly embedded across transcript contexts, with strong enrichment in lncRNAs and intronic regions of protein-coding genes^58–60^. These compartments provide relatively permissive environments in which sequence constraints are reduced and transposable element–derived material can accumulate and be co-opted^59,60^. In contrast, integration within protein-coding exons is comparatively rare, reflecting strong selective pressure to preserve open reading frames and protein function^59^.

Notably, when conserved HERV-derived protein domains are retained within exonic regions, they are not randomly distributed but preferentially localize to transcript termini, particularly last exons and, in protein-coding genes, 3′ UTRs. This positional bias suggests that terminal transcript regions provide a permissive context for the retention of conserved coding remnants without disrupting core coding potential.

The functional relevance of this bias becomes apparent when considering the regulatory landscape of transcript termini. The 3′ UTR is a central hub for post-transcriptional control, coordinating mRNA stability, localization, and translational efficiency^61,62^. The enrichment of conserved HERV-derived sequences in these regions therefore places them within transcript domains that are intrinsically suited to modulate RNA behavior.

In addition, the retention of conserved retroviral domains within terminal exons suggests that remnants of ancestral coding capacity may be functionally repurposed. Beyond contributing to RNA-level regulation, these sequences may give rise to short open reading frames (sORFs) compatible with transcript termini, raising the possibility of translation into micropeptides. Ribosome profiling studies have highlighted the widespread production of such peptides from non-canonical ORFs, particularly within non-coding transcripts^63^.

Consistent with this view, our analyses indicate that candidate HERV-derived micropeptides are more frequently associated with lncRNA contexts than with 3′ UTRs or other regions of protein-coding genes. This pattern suggests that lncRNAs provide a particularly permissive substrate for the emergence of novel coding potential, whereas terminal exons of protein-coding genes represent a more constrained but still functionally relevant environment. In this context, HERV-derived sequences may provide a rich substrate for the emergence of novel micropeptides, linking their evolutionary origin to previously underappreciated layers of translational output.

A particularly striking implication of our results is the emergence of a subclass of lncRNAs that are largely or entirely composed of HERV-derived sequences. Although thousands of lncRNAs overlap HERV internal regions, functional annotation for these transcripts is remarkably sparse, especially among those with the highest HERV content. This observation highlights a fundamental interpretative challenge: whether HERV-dominated lncRNAs represent functional regulatory molecules that remain largely unexplored, or instead reflect byproducts of pervasive transcription arising from retroviral loci embedded in the genome. A balanced view likely encompasses both possibilities. Some of these transcripts may have been co-opted into bona fide regulatory roles, leveraging the rich repertoire of RNA-associated features encoded by HERV sequences, including RBP interaction sites and RNA secondary structures with the potential to mediate protein binding or dsRNA formation^19,64^. Others may represent transcriptional byproducts with limited intrinsic function. However, such pervasive transcription may not be biologically neutral. Low-level expression of transposable element–derived RNAs has been proposed to contribute to basal activation of innate immune pathways through the generation of dsRNA species, providing a tonic background of immune surveillance^65^. In this context, HERV-dominated lncRNAs—particularly those embedded in genomic configurations compatible with antisense transcription—may represent an important and previously underappreciated source of immunogenic RNA. More broadly, the strong enrichment of RNA regulatory features observed across HERV sequences suggests that even transcripts initially arising as byproducts could provide a substrate for evolutionary innovation. Within this framework, HERV-dominated lncRNAs can be conceptualized as a distinct and largely unexplored class of transcripts, spanning a continuum from non-functional transcriptional output to emergent regulatory and immunomodulatory elements.

A major conceptual advance emerging from this study is the recognition that SPARCS-like configurations^25^ are not rare or exceptional features, but instead represent a widespread genomic architecture embedded across the human transcriptome. By identifying more than 6,500 antisense LTR insertions located within transcript termini, our results substantially expand the scope of previously described SPARCS elements and indicate that the potential for dsRNA formation is broadly encoded at the genomic level. Notably, these loci are enriched in genes associated with immune signaling pathways, suggesting a functional link between antisense LTR insertions at transcript termini and innate immune regulation. Importantly, this dsRNA-forming potential appears to be largely decoupled from classical interferon-driven retroviral activation^66–68^. SPARCS-like LTRs are neither enriched for interferon-responsive transcription factor binding motifs nor preferentially associated with structurally intact retroviral elements, indicating that their regulatory logic differs from that of inducible HERV expression. Instead, these findings support a model in which the capacity to generate dsRNA may arise as an intrinsic property of genomic organization, independent of specific activation states. One intriguing possibility is that such configurations could participate in self-reinforcing regulatory circuits, whereby basal or context-dependent transcription of antisense HERV-derived sequences generates dsRNA that activates innate immune pathways, potentially promoting further expression of immunogenic transcripts. Within this framework, the enrichment of SPARCS-like configurations in lncRNAs is particularly notable. Non-coding transcripts may provide a permissive context in which dsRNA-forming architectures can be tolerated and maintained, potentially acting as reservoirs or buffers of immunogenic RNA. Together, these observations suggest that HERV-derived antisense architectures may constitute a pervasive and previously underappreciated layer of transcriptome organization, linking genomic structure to RNA-mediated immune sensing.

Taken together, our findings support a unified model in which HERV sequences act as RNA regulatory modules operating across multiple layers of transcript biology. At the sequence level, they encode regulatory features such as RBP interaction motifs and RNA structures. At the architectural level, they are incorporated into host transcripts, most commonly within intronic regions, but—with respect to conserved protein domains—preferentially retained in terminal exons and 3′ untranslated regions. At the functional level, these features give rise to diverse outputs, including modulation of RNA processing, potential translation into micropeptides, and activation of innate immune sensing through dsRNA formation. Within this framework, individual HERV loci contribute context-dependent combinations of regulatory properties that collectively shape RNA behavior.

This model has broad implications for understanding the role of transposable elements in human biology and disease. Transposable element–derived RNAs are known to contribute to innate immune activation through dsRNA-mediated viral mimicry^65^. Our results extend this concept by demonstrating that SPARCS-like configurations are widespread across the genome and enriched in immune-related genes, indicating that dsRNA-forming potential may be intrinsically encoded in genomic organization. The pervasive embedding of HERV-derived features within regulatory regions of transcripts suggests that transposable elements can influence RNA processing and gene expression beyond classical transcriptional control. These properties are particularly relevant in pathological contexts characterized by transposable element dysregulation, including cancer^12–14^, neurological disorders^15,16^, and autoimmune disease^17^, where aberrant expression of HERV-derived transcripts may reshape RNA regulatory networks and immune signaling pathways.

Several limitations of this study should be acknowledged. Our analyses are based primarily on sequence-derived predictions and genomic annotations, and therefore do not directly demonstrate functional binding of RBPs, translation of predicted micropeptides, or the formation of dsRNA in vivo. In particular, the identification of SPARCS-like configurations reflects genomic potential for dsRNA formation rather than experimentally validated RNA structures. Accordingly, this work should be interpreted as a systematic and exploratory framework for identifying candidate regulatory features at genome scale. However, the strength of this work lies in its systematic and genome-wide integration of multiple regulatory layers, providing a comprehensive framework for generating testable hypotheses across thousands of loci.

Future studies integrating transcriptomic and epigenomic data across physiological and disease contexts will be essential to define the activity and functional relevance of HERV-derived RNA features at single-cell resolution. Experimental approaches such as CLIP-seq^69^, ribosome profiling^70^, and direct detection of dsRNA^71,72^ will be critical to validate the regulatory roles proposed here. In this context, the resources generated in this study, together with existing platforms such as HERVarium^5^, provide a foundation for the exploration of HERV-derived regulatory elements and their contribution to human biology.

In summary, our results establish HERV sequences as a pervasive and functionally versatile component of the human transcriptome, encoding a previously underappreciated layer of RNA regulatory potential. By linking encoded regulatory features, transcript context, and distinct functional consequences—including RNA regulation, micropeptide production, and dsRNA-mediated immune sensing—this work provides a conceptual framework for understanding how endogenous retroviral elements contribute to the complexity of RNA-mediated regulation in both normal physiology and disease.

## Methods

### Retrieval of HERV internal region and LTR annotations from published resources

HERV internal region sequences, provided in FASTA and BED formats together with associated internal domain annotations, were retrieved from the Zenodo repository *Domain-Level Annotation and Conservation of Human Endogenous Retroviruses in the GRCh38 Genome*^6,73^. HERV LTR coordinates were obtained from the Zenodo repository *Structural Annotation and Transcription Factor Motif Mapping of Human Endogenous Retrovirus LTRs* ^5,74^.

### Definition of HERV transcriptional units and locus identifiers

To facilitate consistent locus-level analyses across genomic, structural, and regulatory annotations, HERV internal regions were extended to define unified locus identifiers by incorporating proximal LTRs. Internal regions were extended to include the nearest upstream and downstream LTRs of the same subfamily and strand within 200 bp, retaining only the closest valid element on each side. Extensions that would have introduced overlaps with neighboring loci were reverted to original boundaries. The resulting loci were assigned unique identifiers and represented in GTF format, enabling consistent tracking of individual HERV insertions across all analyses. These locus-level identifiers were used as a unifying framework for integrating domain annotations, regulatory features, and transcript-context overlaps, whereas LTR-specific features were incorporated using external mapping resources.

### Overview of the analysis workflow

The analysis pipeline consisted of three main stages. First, genomic, structural, and regulatory features of HERV internal regions and LTRs were systematically annotated. Particularly RBPBS, and transcript context. Second, these annotations were integrated into a unified locus-level table capturing structural, regulatory, and genomic context information for each HERV internal region. Third, transcript-level and gene-level analyses were performed to characterize the embedding of HERV sequences and conserved domains within host transcripts, including their distribution across transcript features, contribution to lncRNA architecture, and association with regulatory and functional annotations.

### Prediction of RBPBS

Potential RBPBS were predicted by scanning HERV internal region sequences with position weight matrices derived from the ATtRACT database^75^. Human RBP motifs were extracted from ATtRACT and converted to MEME format^76^, retaining matrix identifiers and associated gene annotations. Motifs were restricted to *Homo sapiens* and further filtered to include CisBP-RNA-derived entries^77^ using a custom preprocessing pipeline.

FASTA sequences of HERV internal regions were used as input for motif scanning. Background nucleotide frequencies were estimated directly from the input sequences using a first-order Markov model computed with the MEME suite tool fasta-get-markov, under strand-specific conditions to match the RNA-oriented scanning strategy.

Motif scanning was performed with FIMO^78^ using a p-value threshold of 1 × 10^−4^ and a maximum of 10^6^ stored matches. Scans were conducted in a strand-specific manner (--norc), treating sequences as transcript-oriented and avoiding reverse-complement scoring. FIMO output was parsed to recover genomic coordinates of motif occurrences from sequence identifiers encoding chromosomal position and strand. Coordinates were converted from sequence-relative positions (1-based inclusive) to genomic BED coordinates (0-based, end-exclusive), accounting for sequence orientation. The resulting dataset comprised genome-wide maps of RBPBS across HERV internal regions, including motif identity, genomic coordinates, strand, and statistical significance metrics.

### Annotation of HERV genomic and transcript context

The genomic and transcript context of HERV internal regions, LTRs, and associated protein domains was determined through systematic overlap with GENCODE v48 annotations^79^. Gene, transcript, exon, coding sequence (CDS), and untranslated region (UTR) features were extracted from the GENCODE GTF file and converted to BED format.

Genomic overlaps between HERV internal regions, retroviral protein domains, LTR elements, and transcript features were computed using a standardized bedtools-based pipeline^80^ operating on BED files sorted according to GRCh38 chromosome order.

To generate non-redundant gene-level features, exons were merged across transcripts belonging to the same gene, and introns were defined as the complement of merged exons within gene bodies using bedtools subtract. Overlaps between HERV internal regions and genomic features (genes, exons, introns, CDS, and UTRs) were computed using bedtools intersect in both strand-independent and strand-specific modes. Strand-specific intersections were used to identify HERV insertions co-oriented with host transcripts, which is particularly relevant for transcript incorporation analyses.

For retroviral protein domains, overlaps with transcript-resolved exon, CDS, and UTR annotations were computed using strand-aware base-pair-resolution intersections (bedtools intersect -wo). Overlapping intervals were merged prior to quantification to avoid double counting and to derive unique overlap lengths. Domain–feature overlap fractions were then calculated relative to total domain length.

UTRs were subdivided into 5′ and 3′ components in a transcript-aware manner using CDS boundaries and transcript strand, with CDS start and end coordinates used to assign UTR segments according to transcript orientation. Terminal exons were defined from transcript-resolved exon coordinates using strand-aware exon ordering.

For LTR elements, transcript-feature overlaps were computed in a strand-aware manner to identify antisense insertions relative to host transcripts, enabling detection of genomic configurations compatible with dsRNA formation. Proximity relationships between HERV internal regions and annotated genes or transcription start sites were computed using bedtools closest, both with and without strand constraints.

### Construction of an integrated annotation table for HERV internal regions

To enable downstream analyses, a comprehensive integrated annotation table was constructed for all HERV internal regions by combining genomic, structural, regulatory, and transcript-context features into a unified framework.

For each internal region, retroviral protein domain annotations were aggregated at the locus level, retaining domain identity, domain class (e.g., GAG, POL, ENV), and conservation metrics. LTR-associated information was incorporated by linking each internal region to its corresponding 5′ and 3′ LTRs using the file HERV_internal_LTR_map.tsv from the Zenodo repository *Structural Annotation and Transcription Factor Motif Mapping of Human Endogenous Retrovirus LTRs*^5,74^. Integrated LTR features included LTR length, transcription factor binding motif burden, and reconstruction confidence.

RBPBS information was summarized per locus by aggregating motif occurrences identified by FIMO, including both total motif burden and the number of unique RBPs associated with each internal region. Genomic context annotations were integrated from overlap and proximity analyses, including compact summaries of overlapping genomic features and associated gene biotypes. The final integrated table therefore captured, for each HERV internal region, its structural features, regulatory potential, and genomic context, and included the locus-level identifiers defined above to enable consistent tracking across analyses. This table served as the main input for downstream transcript-level and gene-level analyses.

### Summarization of transcript context of conserved HERV domains

To characterize the transcript-level embedding of conserved HERV protein domains, we summarized domain–transcript overlap annotations derived from the previous step. Input tables contained, for each conserved domain instance, its overlap with exon, CDS, and UTR features, together with positional information including terminal exon status. Analyses were restricted to domain–transcript associations satisfying predefined conservation and overlap criteria. Specifically, domains were required to exhibit ≥40% coverage of the HMM profile and ≥80% overlap of their genomic length with transcript features.

Transcripts were first stratified by gene biotype into protein-coding and lncRNA categories. Within each category, conserved exon-overlapping domains were classified according to whether they occurred in terminal or non-terminal exons, and the proportion in each class was computed to assess enrichment in transcript termini.

For protein-coding transcripts, conserved domains located in terminal exons were further classified according to their embedding within CDS, 5′ UTR, or 3′ UTR regions. This allowed assessment of whether terminal exon-associated domains preferentially reside in coding or non-coding portions of transcripts.

Finally, the composition of retroviral domain classes retained in terminal exons was quantified separately for lncRNA and protein-coding transcripts. Domain classes were aggregated and the number of unique conserved domain instances per class was computed for each transcript category.

### Quantification of HERV contribution to lncRNA transcripts

The contribution of HERV internal regions to lncRNA architecture was quantified by integrating genomic overlap information with transcript annotation data. HERV internal regions overlapping lncRNA genes on the same strand were first identified from genome-wide intersection analyses. For each lncRNA gene, all overlapping HERV loci were collected.

To enable accurate base-pair–level quantification of HERV contribution, gene body and exon intervals were reconstructed directly from GENCODE v48 annotations. Gene body regions were defined using full gene span coordinates, and exon intervals were merged per gene to generate non-redundant exon representations.

HERV intervals were intersected with these reconstructed gene and exon intervals, and overlapping segments were merged to avoid double counting. For each gene, the total number of base pairs contributed by HERV insertions within the gene body and within exons was calculated and normalized by total gene body length and total exon length, respectively.

Based on the fraction of lncRNA gene body overlapped by HERV internal sequence, transcripts were classified into four categories: minor (≤10%), substantial (>10–50%), dominant (>50–80%), and predominant (>80%). This framework enabled stratification of lncRNAs according to the degree of retroviral sequence incorporation.

### Functional annotation and literature mining of HERV-associated lncRNAs

To characterize the functional landscape of HERV-associated lncRNAs, annotations from curated databases and large-scale literature mining were integrated.

Functional annotations were retrieved from LncRNAWiki^81^. Genes were matched using a hierarchical strategy based on Ensembl gene identifiers, transcript identifiers, gene symbols, and known synonyms, with normalization steps applied to harmonize naming conventions. Matched annotations were aggregated at the gene level to summarize reported functional mechanisms, biological processes, molecular functions, pathway associations, and disease links. In parallel, GO annotations^48,49^ were retrieved from Ensembl BioMart^82,83^ and summarized per gene.

To complement curated resources, functional evidence was extracted directly from the biomedical literature through large-scale text mining. PubMed abstracts were retrieved via the NCBI Entrez API^84^ using lncRNA-related queries. Abstracts published from 2000 onward were downloaded together with associated metadata, including PMID, year, and title.

Natural language processing was performed using spaCy^85^. Candidate lncRNA mentions were detected using regular expression patterns capturing common naming conventions, including LINC identifiers, antisense transcript names, and Ensembl gene IDs. Candidate names were normalized and filtered to remove ambiguous or non-informative terms.

For each sentence containing candidate lncRNA mentions, functional context was inferred through predefined keyword sets capturing mechanistic, phenotypic, associative, pathway-related, and regulatory terms. Sentences were classified into categories such as mechanistic evidence, perturbation-based functional evidence, associative observations, or list-like descriptions. A tiered evidence score was assigned, prioritizing mechanistic and perturbation-based statements over descriptive or associative evidence.

Only informative sentences containing both candidate lncRNA mentions and relevant functional terms were retained. For each lncRNA–function association, structured information including functional keywords, sentence class, evidence tier, and the original sentence was extracted. Results were aggregated across abstracts to generate a gene-level dataset of literature-derived functional evidence.

Finally, LncRNAWiki, GO, and literature-derived annotations were integrated into a unified framework, enabling classification of HERV-associated lncRNAs according to the presence and type of functional support.

### Identification of SPARCS-like antisense LTR configurations

SPARCS-like configurations were identified by integrating HERV LTR annotations with transcript-context information derived from GENCODE. Candidate loci were defined as LTR insertions occurring in antisense orientation relative to host transcripts and located within transcript termini, specifically within 3′ UTRs of protein-coding genes or within terminal exons of lncRNAs^25^. To avoid redundancy arising from multiple transcript isoforms, candidates were collapsed to unique LTR instances based on genomic coordinates while preserving aggregated information on associated genes and transcripts.

The resulting SPARCS-like set was further integrated with LTR structural annotations derived from U3–R–U5 segmentation analyses retrieved from the Zenodo repository *Structural Annotation and Transcription Factor Motif Mapping of Human Endogenous Retrovirus LTRs*^5,74^. For each LTR, incorporated structural features included the presence of reconstructed U3 and U5 regions, and overall annotation confidence. This enabled comparison of SPARCS-like LTRs against background LTR sets, including all annotated LTRs and solo LTRs.

Differences in structural integrity and annotation confidence between SPARCS-like and background LTRs were evaluated statistically. Fisher’s exact tests were used for categorical comparisons, including proportions of successfully reconstructed elements, and Wilcoxon rank-sum tests were used for continuous measures such as confidence scores.

### Analysis of interferon-responsive regulatory features in LTRs

To characterize interferon-related regulatory potential in HERV LTRs, transcription factor binding motifs were analyzed using previously computed FIMO scans across all annotated LTR sequences, retrieved from the Zenodo repository *Structural Annotation and Transcription Factor Motif Mapping of Human Endogenous Retrovirus LTRs*^5,74^. Analyses focused on STAT1::STAT2 heterodimer motifs, representing type I/III interferon signaling, STAT1 motifs, representing type II interferon signaling, and IRF family motifs^50–52^.

To obtain robust motif counts, overlapping and redundant hits within each LTR were collapsed using a minimum overlap threshold of 50%, ensuring that each predicted binding site was represented only once. For each LTR, motif burden was quantified as the total number of unique sites, best motif scores and p-values, and motif density normalized by LTR length.

Local motif clustering was assessed by scanning each LTR with 75-bp sliding windows. This was performed independently for STAT and IRF motifs and jointly to identify co-localized regions. For each LTR, the densest window of combined STAT and IRF motifs was recorded together with the number of STAT and IRF sites it contained.

LTRs were then assigned to categories reflecting their predicted interferon-responsive potential based on local motif architecture. Motif organization was evaluated within sliding windows of 75 bp, and classification was based on the composition of the densest window identified for each LTR. LTRs containing at least one STAT motif and one IRF motif within the same window were classified as having synergistic potential and further subdivided into very high (≥3 total motifs within the window) and high (exactly one STAT and one IRF motif within the window) categories. LTRs exhibiting strong clustering of a single motif class (≥3 STAT or ≥3 IRF motifs within a window) were also classified as high. LTRs with exactly two motif occurrences of the same class within the window were classified as medium, whereas those with a single motif occurrence were classified as low. This classification provided a proxy for the presence of cooperative regulatory modules consistent with interferon-responsive enhancers^50,52,86^..

SPARCS-like LTRs were intersected with interferon motif annotations to assess whether dsRNA-compatible genomic configurations are associated with canonical interferon-responsive regulatory architectures. Enrichment of interferon-associated motif classes was evaluated at the LTR level using Fisher’s exact tests, comparing the proportion of LTRs classified as interferon-responsive between SPARCS-like candidates and background LTR sets. Additional analyses focused on synergistic architectures corresponding to co-localized STAT and IRF motifs.

At the gene level, enrichment analyses were performed for protein-coding genes associated with SPARCS-like LTRs using GO Biological Process annotations. Over-representation analysis was conducted with the clusterProfiler R package^87^ with all protein-coding genes annotated in the GENCODE v48 reference used as the background universe. Statistical significance was assessed using Benjamini–Hochberg multiple-testing correction with an adjusted p-value threshold of 0.05. Complementary analyses were performed using immune-related gene sets from the MSigDB^88^ C7 (ImmuneSigDB) collection, using the enricher function from clusterProfiler with the same correction threshold.

## Data availability

The human reference genome assembly (GRCh38) was obtained from the GENCODE project (release 48; https://ftp.ebi.ac.uk/pub/databases/gencode/Gencode_human/release_48/GRCh38.primary_assembly.genome.fa.gz).

HERV internal region annotations were retrieved from the Zenodo repository *Domain-Level Annotation and Conservation of Human Endogenous Retroviruses in the GRCh38 Genome*^6,73^ (https://doi.org/10.5281/zenodo.17662456), including the files HERV_internal_v5.bed, HERV_internal_sequences_v5.fasta, HERV_domains_v5.bed, and HERV_loci_annotated_domains_v5.tsv.

HERV LTR annotations were obtained from the Zenodo repository *Structural Annotation and Transcription Factor Motif Mapping of Human Endogenous Retrovirus LTRs*^5,74^ (https://doi.org/10.5281/zenodo.19593457), including the files HERV_ltr.bed, HERV_internal_LTR_map.tsv, HERV_LTR_U3_R_U5_catalogue.tsv, and HERV_LTR_fimo_results_tsv.tar.xz.

RBPBS motifs were obtained from the ATtRACT database (https://attract.cnic.es; ^75^), available at https://attract.cnic.es/attract/static/ATtRACT.zip.

All HERV annotations generated in this study, together with their description, are publicly available in Zenodo (https://doi.org/10.5281/zenodo.19661036).

## Code availability

All code used for data processing and analysis in this study is available as a collection of Python, Bash, and R scripts at https://github.com/funcgen/herv-rna-regulation.

## Competing interest statement

The authors declare no competing interests.

## Acknowledgements

We acknowledge the use of artificial intelligence tools (ChatGPT, OpenAI) to assist in code drafting and language editing.

Author contributions: T.M-A and A.E-C conceived the project. T.M-A performed the bioinformatic analyses. T.M-A and A.E-C wrote the manuscript. A.E-C supervised the project. A.P. contributed to data review, scientific discussions and provided critical feedback on the manuscript.

## Funding

This publication and all its results are supported by the AGAUR-FI predoctoral grant program (2025 FI-1 00642) Joan Oró, from the Secretariat for Universities and Research of the Department of Research and Universities of the Government of Catalonia, and by the European Social Fund Plus.

AP has received funding from the Secretariat for Universities and Research of the Ministry of Business and Knowledge of the Government of Catalonia (2021SGR00899), and the Instituto de Salud Carlos III (ISCIII) (FIS PI23/00835), ‘Fondo Europeo de Desarrollo Regional (FEDER), Unión Europea, una manera de hacer Europa’. Institutional support to CNAG was from the Spanish Ministry of Science, Innovation and Universities, Fondo de Investigaciones Sanitarias cofunded with ERDF funds (PI19/01772), the 2014–2020 Smart Growth Operating Program, and the Generalitat de Catalunya through the Departament de Recerca i Universitats and Departament de Salut.

## Notes

https://doi.org/10.5281/zenodo.19661036

https://github.com/funcgen/herv-rna-regulation

### Competing Interest Statement

The authors have declared no competing interest.

## References

1. Cordaux, R. & Batzer, M. A. The impact of retrotransposons on human genome evolution. Nat. Rev. Genet. 10, 691–703 (2009).

2. Du, A. Y., Chobirko, J. D., Zhuo, X., Feschotte, C. & Wang, T. Regulatory transposable elements in the encyclopedia of DNA elements. Nat. Commun. 15, 7594 (2024).

3. Griffiths, D. J. Endogenous retroviruses in the human genome sequence. Genome Biol. 2, REVIEWS1017 (2001).

4. Ito, J. et al. Systematic identification and characterization of regulatory elements derived from human endogenous retroviruses. PLOS Genet. 13, e1006883 (2017).

5. Montserrat-Ayuso, T., Pujol, A. & Esteve-Codina, A. Regulatory Features and Functional Specialization of Human Endogenous Retroviral LTRs: A Genome-Wide Annotation and Analysis via HERVarium. 2026.02.17.706328 Preprint at 10.64898/2026.02.17.706328 (2026).

6. Montserrat-Ayuso, T., Pujol, A. & Esteve-Codina, A. A comprehensive annotation of conserved protein domains in human endogenous retroviruses. NAR Genomics Bioinforma. 8, lqag013 (2026).

7. Bannert, N. & Kurth, R. The evolutionary dynamics of human endogenous retroviral families. Annu. Rev. Genomics Hum. Genet. 7, 149–173 (2006).

8. Wang, J., Lu, X., Zhang, W. & Liu, G.-H. Endogenous retroviruses in development and health. Trends Microbiol. 32, 342–354 (2024).

9. Grandi, N. & Tramontano, E. Human Endogenous Retroviruses Are Ancient Acquired Elements Still Shaping Innate Immune Responses. Front. Immunol. 9, (2018).

10. Mi, S. et al. Syncytin is a captive retroviral envelope protein involved in human placental morphogenesis. Nature 403, 785–789 (2000).

11. Muir, A., Lever, A. & Moffett, A. Expression and Functions of Human Endogenous Retroviruses in the Placenta: An Update. Placenta 25, S16–S25 (2004).

12. Li, Z. et al. Expression of HERV-K correlates with status of MEK-ERK and p16INK4A-CDK4 pathways in melanoma cells. Cancer Invest. 28, 1031–1037 (2010).

13. Zhao, J. et al. Expression of Human Endogenous Retrovirus Type K Envelope Protein is a Novel Candidate Prognostic Marker for Human Breast Cancer. Genes Cancer 2, 914–922 (2011).

14. Rycaj, K. et al. Cytotoxicity of human endogenous retrovirus K-specific T cells toward autologous ovarian cancer cells. Clin. Cancer Res. Off. J. Am. Assoc. Cancer Res. 21, 471–483 (2015).

15. Küry, P. et al. Human Endogenous Retroviruses in Neurological Diseases. Trends Mol. Med. 24, 379–394 (2018).

16. Duarte, R. R. R., Nixon, D. F. & Powell, T. R. Ancient viral DNA in the human genome linked to neurodegenerative diseases. Brain. Behav. Immun. 123, 765–770 (2025).

17. Viret, C. & Bynoe, M. S. Human Endogenous Retroviruses Expression in Autoimmunity. Yale J. Biol. Med. 97, 521–528 (2024).

18. Foroushani, A. K. et al. Posttranscriptional regulation of human endogenous retroviruses by RNA-binding motif protein 4, RBM4. Proc. Natl. Acad. Sci. 117, 26520–26530 (2020).

19. Onoguchi, M., Zeng, C., Matsumaru, A. & Hamada, M. Binding patterns of RNA-binding proteins to repeat-derived RNA sequences reveal putative functional RNA elements. NAR Genomics Bioinforma. 3, lqab055 (2021).

20. Pe, K. & Dl, M. A human endogenous retrovirus suppresses translation of an associated fusion transcript, PLA2L. J. Virol. 72, (1998).

21. Kitao, K., Tanikaga, T. & Miyazawa, T. Identification of a post-transcriptional regulatory element in the human endogenous retroviral syncytin-1. J. Gen. Virol. 100, 662–668 (2019).

22. Gebrie, A. Transposable elements as essential elements in the control of gene expression. Mob. DNA 14, 9 (2023).

23. Wang, Y. et al. Endogenous miRNA sponge lincRNA-RoR regulates Oct4, Nanog, and Sox2 in human embryonic stem cell self-renewal. Dev. Cell 25, 69–80 (2013).

24. Kulski, J. K. Long Noncoding RNA HCP5, a Hybrid HLA Class I Endogenous Retroviral Gene: Structure, Expression, and Disease Associations. Cells 8, 480 (2019).

25. Cañadas, I. et al. Tumor innate immunity primed by specific interferon-stimulated endogenous retroviruses. Nat. Med. 24, 1143–1150 (2018).

26. Jansz, N. & Faulkner, G. J. Endogenous retroviruses in the origins and treatment of cancer. Genome Biol. 22, 147 (2021).

27. Zhang, J. et al. LIN28 Regulates Stem Cell Metabolism and Conversion to Primed Pluripotency. Cell Stem Cell 19, 66–80 (2016).

28. Wang, W. et al. Profiling the role of m6A effectors in the regulation of pluripotent reprogramming. Hum. Genomics 18, 33 (2024).

29. Liu, H.-L. et al. The role of RNA splicing factor PTBP1 in neuronal development. Biochim. Biophys. Acta BBA - Mol. Cell Res. 1870, 119506 (2023).

30. Bugg, D. et al. MBNL1 drives dynamic transitions between fibroblasts and myofibroblasts in cardiac wound healing. Cell Stem Cell 29, 419–433.e10 (2022).

31. Saito, Y. et al. Differential NOVA2-mediated splicing in excitatory and inhibitory neurons regulates cortical development and cerebellar function. Neuron 101, 707–720.e5 (2019).

32. Choudhury, R. et al. The splicing activator DAZAP1 integrates splicing control into MEK/Erk regulated cell proliferation and migration. Nat. Commun. 5, 3078 (2014).

33. Li, J. et al. HNRNPK maintains epidermal progenitor function through transcription of proliferation genes and degrading differentiation promoting mRNAs. Nat. Commun. 10, 4198 (2019).

34. Berto, S., Usui, N., Konopka, G. & Fogel, B. L. ELAVL2-regulated transcriptional and splicing networks in human neurons link neurodevelopment and autism. Hum. Mol. Genet. 25, 2451–2464 (2016).

35. Haemmerle, M. W. et al. RNA binding proteins PCBP1 and PCBP2 regulate pancreatic β cell translation. Mol. Metab. 98, 102175 (2025).

36. He, X., Yuan, J. & Wang, Y. G3BP1 binds to guanine quadruplexes in mRNAs to modulate their stabilities. Nucleic Acids Res. 49, 11323–11336 (2021).

37. Zdradzinski, M. D. et al. KHSRP-mediated decay of axonally localized prenyl-Cdc42 mRNA slows nerve regeneration. PLoS Genet. 21, e1011916 (2025).

38. Shu, H. et al. FMRP links optimal codons to mRNA stability in neurons. Proc. Natl. Acad. Sci. 117, 30400–30411 (2020).

39. George, J. et al. RNA-binding protein FXR1 drives cMYC translation by recruiting eIF4F complex to the translation start site. Cell Rep. 37, (2021).

40. Rodrigues, K. S., Petroski, L. P., Utumi, P. H., Ferrasa, A. & Herai, R. H. IARA: a complete and curated atlas of the biogenesis of spliceosome machinery during RNA splicing. Life Sci. Alliance 6, (2023).

41. Geuens, T., Bouhy, D. & Timmerman, V. The hnRNP family: insights into their role in health and disease. Hum. Genet. 135, 851–867 (2016).

42. Qi, Y., Wang, M. & Jiang, Q. PABPC1——mRNA stability, protein translation and tumorigenesis. Front. Oncol. 12, (2022).

43. Sanduja, S., Blanco, F. F. & Dixon, D. A. The roles of TTP and BRF proteins in regulated mRNA decay. Wiley Interdiscip. Rev. RNA 2, 42–57 (2010).

44. Forouzanfar, M. et al. Intracellular functions of RNA-binding protein, Musashi1, in stem and cancer cells. Stem Cell Res. Ther. 11, 193 (2020).

45. Grifone, R. et al. Rbm24 displays dynamic functions required for myogenic differentiation during muscle regeneration. Sci. Rep. 11, 9423 (2021).

46. Fuchs, N. V. et al. Human endogenous retrovirus K (HML-2) RNA and protein expression is a marker for human embryonic and induced pluripotent stem cells. Retrovirology 10, 115 (2013).

47. Wang, T., Doucet-O’Hare, T. T., Henderson, L., Abrams, R. P. M. & Nath, A. Retroviral Elements in Human Evolution and Neural Development. *J*. Exp. Neurol. 2, 1–9 (2021).

48. Ashburner, M. et al. Gene Ontology: tool for the unification of biology. Nat. Genet. 25, 25–29 (2000).

49. The Gene Ontology Consortium. The Gene Ontology knowledgebase in 2026. Nucleic Acids Res. 54, D1779–D1792 (2026).

50. Michalska, A., Blaszczyk, K., Wesoly, J. & Bluyssen, H. A. R. A Positive Feedback Amplifier Circuit That Regulates Interferon (IFN)-Stimulated Gene Expression and Controls Type I and Type II IFN Responses. Front. Immunol. 9, (2018).

51. Au-Yeung, N., Mandhana, R. & Horvath, C. M. Transcriptional regulation by STAT1 and STAT2 in the interferon JAK-STAT pathway. JAK-STAT 2, e23931 (2013).

52. Mogensen, T. H. IRF and STAT Transcription Factors - From Basic Biology to Roles in Infection, Protective Immunity, and Primary Immunodeficiencies. Front. Immunol. 9, (2019).

53. Izsvák, Z., Ma, J., Singh, M. & Hurst, L. D. Co-option of an endogenous retrovirus (LTR7-HERVH) in early human embryogenesis: becoming useful and going unnoticed. Mob. DNA 16, 27 (2025).

54. Göke, J. et al. Dynamic transcription of distinct classes of endogenous retroviral elements marks specific populations of early human embryonic cells. Cell Stem Cell 16, 135–141 (2015).

55. Ohnuki, M. et al. Dynamic regulation of human endogenous retroviruses mediates factor-induced reprogramming and differentiation potential. Proc. Natl. Acad. Sci. 111, 12426–12431 (2014).

56. Zhang, Y. et al. Transcriptionally active HERV-H retrotransposons demarcate topologically associating domains in human pluripotent stem cells. Nat. Genet. 51, 1380–1388 (2019).

57. Grow, E. J. et al. Intrinsic retroviral reactivation in human preimplantation embryos and pluripotent cells. Nature 522, 221–225 (2015).

58. Göke, J. & Ng, H. H. CTRL+INSERT: retrotransposons and their contribution to regulation and innovation of the transcriptome. EMBO Rep. 17, 1131–1144 (2016).

59. Kapusta, A. et al. Transposable elements are major contributors to the origin, diversification, and regulation of vertebrate long noncoding RNAs. PLoS Genet. 9, e1003470 (2013).

60. Drongitis, D., Aniello, F., Fucci, L. & Donizetti, A. Roles of Transposable Elements in the Different Layers of Gene Expression Regulation. Int. J. Mol. Sci. 20, 5755 (2019).

61. Hong, D. & Jeong, S. 3’UTR Diversity: Expanding Repertoire of RNA Alterations in Human mRNAs. Mol. Cells 46, 48–56 (2023).

62. Mayya, V. K. & Duchaine, T. F. Ciphers and Executioners: How 3′-Untranslated Regions Determine the Fate of Messenger RNAs. Front. Genet. 10, (2019).

63. Pan, J. et al. Functional Micropeptides Encoded by Long Non-Coding RNAs: A Comprehensive Review. Front. Mol. Biosci. 9, (2022).

64. Sadeq, S., Al-Hashimi, S., Cusack, C. M. & Werner, A. Endogenous Double-Stranded RNA.Non-Coding RNA 7, 15 (2021).

65. Kassiotis, G. & Stoye, J. P. Immune responses to endogenous retroelements: taking the bad with the good. Nat. Rev. Immunol. 16, 207–219 (2016).

66. Manghera, M., Ferguson-Parry, J., Lin, R. & Douville, R. N. NF-κB and IRF1 Induce Endogenous Retrovirus K Expression via Interferon-Stimulated Response Elements in Its 5′ Long Terminal Repeat. J. Virol. 90, 9338–9349 (2016).

67. Russ, E., Mikhalkevich, N. & Iordanskiy, S. Expression of Human Endogenous Retrovirus Group K (HERV-K) HML-2 Correlates with Immune Activation of Macrophages and Type I Interferon Response. Microbiol. Spectr. 11, e04438–22 (2023).

68. Le, E. et al. Type I interferons increase expression of endogenous retrovirus K102 and envelope protein in myeloid cells from patients with autoimmune disease. Mob. DNA 16, 36 (2025).

69. Lee, F. C. Y. & Ule, J. Advances in CLIP Technologies for Studies of Protein-RNA Interactions. Mol. Cell 69, 354–369 (2018).

70. Ingolia, N. T., Ghaemmaghami, S., Newman, J. R. S. & Weissman, J. S. Genome-Wide Analysis in Vivo of Translation with Nucleotide Resolution Using Ribosome Profiling. Science 324, 218–223 (2009).

71. Schonborn, J. et al. Monoclonal antibodies to double-stranded RNA as probes of RNA structure in crude nucleic acid extracts. Nucleic Acids Res. 19, 2993–3000 (1991).

72. Decker, C. J. et al. dsRNA-Seq: Identification of Viral Infection by Purifying and Sequencing dsRNA. Viruses 11, 943 (2019).

73. Montserrat-Ayuso, T., Pujol, A. & Esteve-Codina, A. Domain-Level Annotation and Conservation of Human Endogenous Retroviruses in the GRCh38 Genome. Zenodo 10.5281/zenodo.17662456 (2025).

74. Montserrat-Ayuso, T., Pujol, A. & Esteve-Codina, A. Structural Annotation and Transcription Factor Motif Mapping of Human Endogenous Retrovirus LTRs. Zenodo 10.5281/zenodo.18554805 (2026).

75. Giudice, G., Sánchez-Cabo, F., Torroja, C. & Lara-Pezzi, E. ATtRACT-a database of RNA-binding proteins and associated motifs. Database J. Biol. Databases Curation 2016, baw035 (2016).

76. Bailey, T. L., Johnson, J., Grant, C. E. & Noble, W. S. The MEME Suite. Nucleic Acids Res. 43, W39–W49 (2015).

77. Ray, D. et al. A compendium of RNA-binding motifs for decoding gene regulation. Nature 499, 172–177 (2013).

78. Grant, C. E., Bailey, T. L. & Noble, W. S. FIMO: scanning for occurrences of a given motif. Bioinformatics 27, 1017–1018 (2011).

79. Mudge, J. M. et al. GENCODE 2025: reference gene annotation for human and mouse. Nucleic Acids Res. 53, D966–D975 (2025).

80. Quinlan, A. R. & Hall, I. M. BEDTools: a flexible suite of utilities for comparing genomic features. Bioinformatics 26, 841–842 (2010).

81. Liu, L. et al. LncRNAWiki 2.0: a knowledgebase of human long non-coding RNAs with enhanced curation model and database system. Nucleic Acids Res. 50, D190–D195 (2022).

82. Dyer, S. C. et al. Ensembl 2025. Nucleic Acids Res. 53, D948–D957 (2025).

83. Kinsella, R. J. et al. Ensembl BioMarts: a hub for data retrieval across taxonomic space. Database J. Biol. Databases Curation 2011, bar030 (2011).

84. Sayers, E. W. et al. Database resources of the National Center for Biotechnology Information in 2025. Nucleic Acids Res. 53, D20–D29 (2025).

85. Montani, I., et al. explosion/spaCy v2.2.4.dev0. Zenodo 10.5281/zenodo.3701227 (2020).

86. Gotea, V. et al. Homotypic clusters of transcription factor binding sites are a key component of human promoters and enhancers. Genome Res. 20, 565–577 (2010).

87. Yu, G., Wang, L.-G., Han, Y. & He, Q.-Y. clusterProfiler: an R Package for Comparing Biological Themes Among Gene Clusters. OMICS J. Integr. Biol. 16, 284–287 (2012).

88. Liberzon, A. et al. Molecular signatures database (MSigDB) 3.0. Bioinformatics 27, 1739–1740 (2011).

