## Supplemental Figure for "HERVs as building blocks of RNA regulatory architecture in the human genome"

A

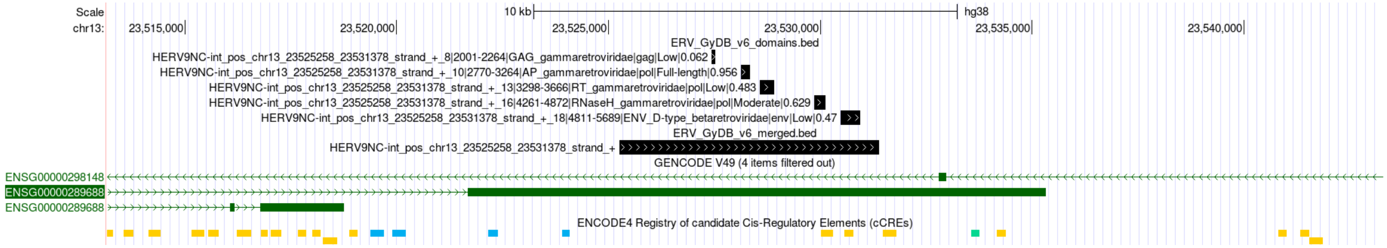


B

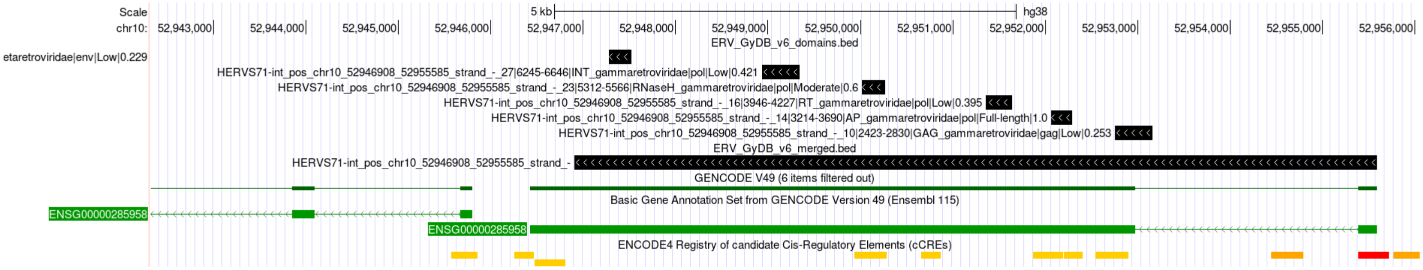


**Supplemental Figure 1. Conserved HERV-derived domains in lncRNA loci and regulatory context.** (A–B) Genome-browser views of representative lncRNA loci harboring conserved HERV-derived domains within terminal exons. Tracks show HERV internal regions, annotated retroviral protein domains, GENCODE gene models, and ENCODE candidate cCREs. In both examples, conserved domains (including protease, RNase H, reverse transcriptase, and Env) are embedded within transcriptionally active regions and coincide with multiple cCREs, indicating integration of retroviral coding remnants within regulatory genomic environments.

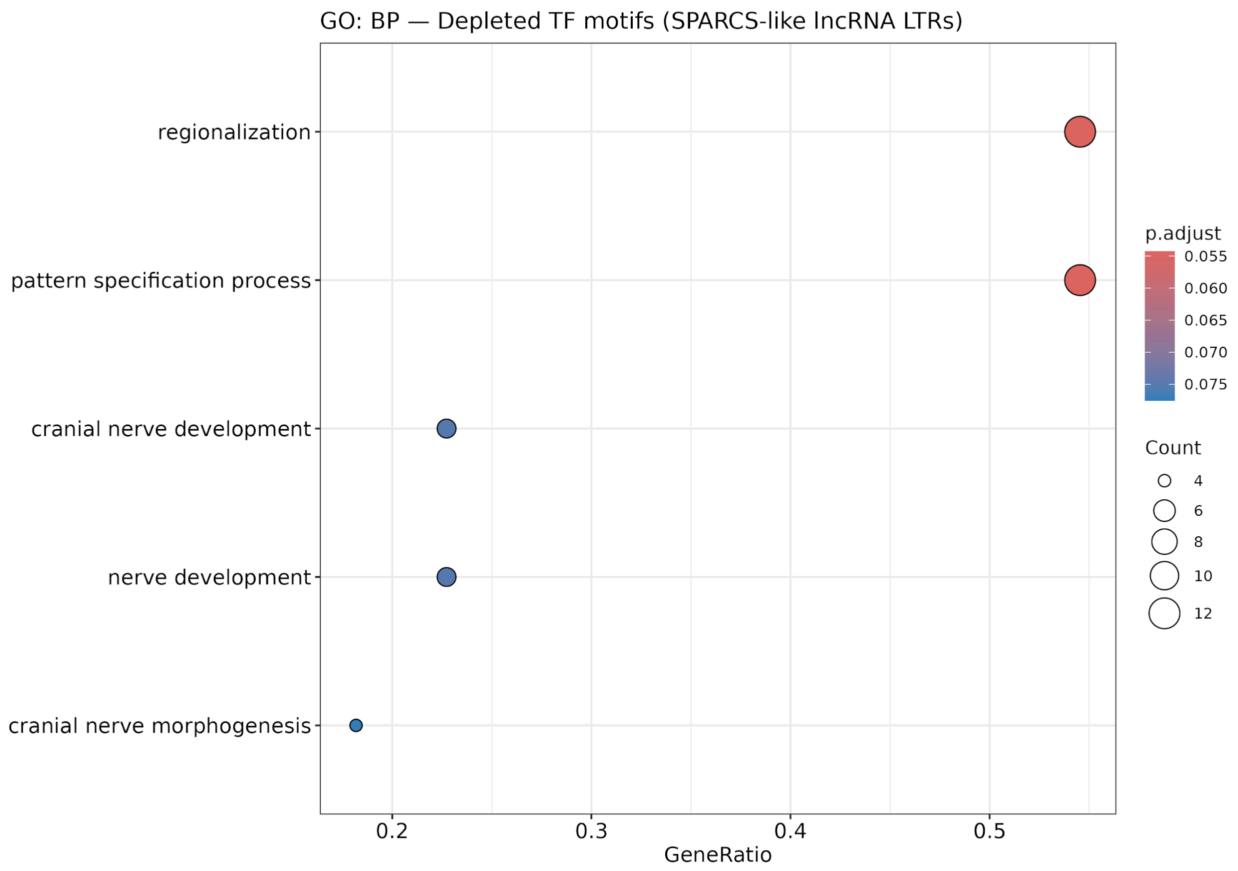


**Supplemental Figure 2. Supplemental Figure 2. Regulatory landscape of SPARCS-like LTRs.** Gene Ontology (GO) enrichment analysis of transcription factors depleted in SPARCS-like LTRs located within lncRNA transcripts. Enriched terms are predominantly associated with developmental processes, particularly regionalization and pattern specification. Dot size represents gene count and color indicates adjusted p-value.
